# First detection of *Ixodiphagus hookeri* (Hymenoptera: Encyrtidae) in *Ixodes ricinus* ticks (Acari: Ixodidae) from multiple locations of Hungary

**DOI:** 10.1101/2022.09.05.506604

**Authors:** Adrienn Gréta Tóth, Róbert Farkas, Mónika Gyurkovszky, Eszter Krikó, Norbert Solymosi

## Abstract

The parasiotid wasp, *Ixodiphagus hookeri* (Hymenoptera: Encyrtidae) is the natural enemy of a wide range of hard and soft tick species. While these encyrtid wasps are supposed to be distributed worldwide, only few studies report about its actual appearance patterns around the globe. Within a shotgun sequencing based metagenome analysis, the occurrence of *I. hookeri* was screened at multiple *Ixodes ricinus* (Acari: Ixodidae) tick sampling points of Hungary, to contribute to the assessment of the appearance patterns of the parasitoid wasps in Central Europe. To our knowledge, the first report of the species in Hungary and the description of the southernmost *I. hookeri* associated geoposition in Central Europe took place within our study. *I. hookeri* infested *I. ricinus* nymphs were detected at five sampling points of Hungary. The results show that the exact distribution range of *I. hookeri* is still barely studied. At the same time, unprecedented public health issues being brought by climate change might require steps towards the exploitation of the tick biocontrol potential or ecological bioindicator role of the parasitoid wasp in the future.

## Introduction

The emergence of vector-borne diseases that can affect both human and animal populations are strongly influenced by the effects of climate change, urbanization and globalization.^1^ Among the most significant arthropod vectors (e.g., ticks, fleas, black flies, mosquitoes, or sand flies) ticks transmit a markedly broad spectrum of pathogenic microorganisms, including various protozoa, rickettsiae, spirochaetes and viruses.^2^ Owing to its geographic position and climatic conditions, Hungary, a landlocked country situated in Central Europe, hosts relatively many, namely at least 27 hard tick species (Ixodidae).^3^ As part of the VectorNet project, the European Centre of Disease Prevention and Control (ECDC) currently monitors seven tick species (*Ixodes ricinus*, *Ixodes persulcatus*, *Dermacentor reticulatus*, *Hyalomma marginatum*, *Rhipicephalus bursa*, *Rhipicephalus sanguineus*, *Ornithodoros* spp.) that commonly transfer diseases to humans and animals.^4^ Naturally, further tick species may also serve as vectors for microorganisms of human or veterinary medical significance.^5^ Out of the seven tick species monitored by ECDC, all except for *Ornithodoros* spp. are hard ticks that are present in Hungary.^3^

Several studies report about the incidence trends of tick-borne encephalitis (TBE) and of Lyme borreliosis (LB) that the most prevalent tick-borne infection in Europe.^6,7^ Enhanced surveillance and diagnostic measures raise awareness of the changing geographical distribution, density, and activity of the *I. ricinus*, the primary vector of TBE and LB in Europe.^8^ As a consequence of climate change, *I. ricinus* tends to appear both at extremes of altitude and latitude apart from its prior range, and its northerly shift within the European continent has also been documented.^9–11^ In case of LB, that has higher country-wise incidence rates than TBE, decade-long trends of incidence rates are not consistent along the countries around the world, increasing and decreasing tendencies both appear. On the other hand, reports of the geographic distribution of LB show a clear expansion, especially towards higher altitudes and latitudes.^6,7^

As climate change accompanied with various sociodemographic alterations brings a time of unprecedented challenges related to vector-borne diseases^12,13^, the need for the development of control methods against tick populations is a public highlight. To address this issue, several methods have been introduced that may focus either on human or animal tick engorgement targets. These control methods often rest on either conventional chemical acaricides that can be encumbered by the resistance of the target pests^14^ or on further alternatives or supplements, such as landscape management, vaccines, pheromone-assisted tools, wild host targeted strategies, genomic control advances, host genotype considerations or biological control methods assisted by the natural enemies of ticks, e.g. entomopathogenic fungi, various species of nematodes or certain parasitoids.^15–19^

A line of the biological control methods against ticks could potentially be the members of Encyrtidae family that are small-sized, parasitoid or hyperparasitoid wasps distributed all around the globe. Due to their efficacy and target specificity, many wasps from this family are used as means of biological pest control, while several additional encyrtid species are documented as promising candidates for this role.^20–22^ *Ixodiphagus* spp., including *Ixodiphagus hookeri*^23^, are encyrtid wasps attacking a wide range of tick species that have received relatively much attention as a specific and effective, natural alternative for biological hard tick control.^24,25^ Interestingly, *I. hookeri* appears to have alternating preferences for the tick species and developmental stage of its hosts at geographically distant locations.^26,27^ In European settings, Ixodiphagus wasps are described to parasitize the larvae and nymphs of hard ticks with a clear predilection for unfed nymphs.^25^ If oviposition occurs to larvae, transstadial transmission through the molting of the ticks to nymphs can also occur.^28^ Wasp eggs start their embryonic development in engorging or engorged nymphs. Wasp larvae feed on tick tissues and emerge as fully grown adults causing the death of the host before it could reach the adult stage.^24,25^

*Ixodiphagus* wasps have been associated with several hard tick genera belonging to the famiIies of Ixodidae and Argasidae, including *Ornithodoros*, *Amblyomma*, *Dermacentor*, *Haemaphysalis*, *Hyalomma*, *Ixodes* and *Rhipicephalus*.^24^ Studies conducted in Europe revealed that *I. ricinus* appears to be the preferred species of the European *I. hookeri*, while another common tick species, *Dermacentor reticulatus* is supposed not to be chosen as a host by the European representatives of the parasitoid wasps.^25^

While *Ixodiphagus* spp. have been detected in many countries and in a diverse range of hard and soft tick species, parasitoid wasps have been less studied in Hungary despite their public health potential by tick-borne diseases. In the present study, our aim was to confirm the presence *I. hookeri* in a diverse set of locations in Hungary using a modern, sensitive metagenomic approach.^29,30^ Due to their high Hungarian prevalence rates, high public health significance and high *Ixodiphagus* wasp hosting potential in Europe, *I. ricinus* ticks were decided to be assessed for the parasitoids. Furthermore, our objective was to contribute to the knowledge collected regarding the life cycle of *I. hookeri* and examine the theory, according to which wasp infested nymphs cannot reach the adult age. Owing to our approach, genomic information of the European populations of *I. hookeri* may also be obtained that can serve as a reference for further studies.

## Materials and Methods

Between March and August of 2019, in a country-wide *I. ricinus* metagenome survey, by flagging and dragging questing ticks were collected from 21 different geolocations in Hungary. The collected ticks were frozen. In the laboratory the ticks were classified taxonomically and 10 nymphs and 10 females of *I. ricinus* were selected per sampling sites (Fig 2). Before DNA extraction the ticks were washed twice by 80% alcohol.

For the DNA isolation the blackPREP Tick DNA/RNA Kit (Analytik Jena GmbH) was used. Isolated total metagenome DNA was used for library preparation. In vitro fragment libraries were prepared using the NEBNext Ultra II DNA Library Prep Kit for Illumina. Paired-end fragment reads were generated on an Illumina NextSeq sequencer using TG NextSeq 500/550 High Output Kit v2 (300 cycles). Primary data analysis (base-calling) was carried out with Bbcl2fastq software (v2.17.1.14, Illumina).

On the raw sequencing data a quality based filtering and trimming was performed by TrimGalore (v.0.6.6, https://github.com/FelixKrueger/TrimGalore), setting 20 as a quality threshold, retaining reads longer than 50 bp only. Using the remained reads a de novo assembly was performed by MEGAHIT (v1.2.9)^31^ using default settings. The resulting contigs were taxonomically classified using Kraken2 (*k* = 35)^32^ with the NCBI non-redundant nucleotide database^33^. Contigs were predicted as *I. hookeri* by taxon classification were checked with BLAST^34^ on the partial sequence of *I. hookeri* Ixo4 28S ribosomal RNA gene (MH077537.1) as a reference. Multiple sequence alignment was done by MAFFT (v7.490)^35^. All data management procedures, analyses and plottings^36,37^ were performed in R environment (v4.2.1).^38^

## Results

Of the 21 female samples examined, we did not find any contigs that could be reasonably assumed to be of *I. hookeri* origin. In five of the 21 nymph samples (b, c, d, g, n), contigs of *I. hookeri* origin were found.

Sequence found in sample b (length: 386, identities: 378/386 (97.9%), no gap):

CTGCTCGACGGTATTCGAATATCACAAGGTGTGTAATCGAGAGCCGCATTTGAATGCGTTCGGCGCGTCGGTCGGCTTCGGCTCGCTCGGTTATTTTACGGACCTGGATGCCGTTACTGGGCGCTTGCCGAGGCTTTTTGTACCGACGACGATCTCGAACCGGCTCTGCGCGCGCGAAAGCGTTCGCGCTCTCGCGCGCTCACCTGTCGGCGACGCTTTTGCTTTGGGTACTTTCAGGACCCGTCTTGAAACACGGACCAAGGAGTCTAACATGTGCGCGAGTCATTGGGTTTTTTTATATTATATTTAAAGCCTAAAGGCGCAATGAAAGTGAAGATACGGCAGGCATTCGTGCCTGAGCCGATCGAGGGAGGATGGCCCGCGTC

Sequence found in sample c (length: 559, identities: 556/559 (99.4%), no gap):

TTCAAGAGTACGTGAAACCGTTCAGGGGTAAACCTGAGAAACCCAAAAGATCGAATGGGGAGATTCAGCGTTCAACGGCCCGTCTGGCTTGCGTGCGACGTCACGATGTCGCGGTGTTATGCGCCCTCGCCGGTGTGTATATACCGTGACACGTCGTCGCTGCGTCATGACCGGCGCCGTCGGCGTGCACTTCTCCCCTAGTAGAACGTCGCGACCCGTTGTGCGTCGGCCAAAGGCTCGAAGGGTAGACTATCGCGCTCTCTCCCCGGAGGGCGCGGCAGACCCTCGAAAGCCCGGCCCGACTGCTCGACGGTATTCGAATATCACAAGGTGTGTAATCGAGAGCCGCATTTGAATGCGTTCGGCGCGTCGGTCGGCTTCGGCTCGCTCGGTTATTTTACGGACCTGGATGCCGTTACTGGGCGCTTGCCGAGGCTTTTTGTACCGACGACGATCTCGAACCGGCTCTGCGCGCGCGAAAGCGTTCGCGCTCTCGCGCGCTCACCTGTCGGCGACGCTTTTGCTTTGGGTACTTTCAGGACCCGTCTTGAAACACGGA

Sequence found in sample d (length: 447, identities: 439/447 (98.2%), no gap):

CTCGAAAGCCCGGCCCGACTGCTCGACGGTATTCGAATATCACAAGGTGTGTAATCGAGAGCCGCATTTGAATGCGTTCGGCGCGTCGGTCGGCTTCGGCTCGCTCGGTTATTTTACGGACCTGGATGCCGTTACTGGGCGCTTGCCGAGGCTTTTTGTACCGACGACGATCTCGAACCGGCTCTGCGCGCGCGAAAGCGTTCGCGCTCTCGCGCGCTCACCTGTCGGCGACGCTTTTGCTTTGGGTACTTTCAGGACCCGTCTTGAAACACGGACCAAGGAGTCTAACATGTGCGCGAGTCATTGGGTTTTTTTATATTATATTTAAAGCCTAAAGGCGCAATGAAAGTGAAGATACGGCAGGCATTCGTGCCTGAGCCGATCGAGGGAGGATGGCCCGCGTCACGATGCGGGCCCGCACTCCCGGGGCGTCTCGCGCTCATTGCG

Sequence found in sample g (length: 607, identities: 445/453 (98.2%), no gap):

AGCCCGGCCCGACTGCTCGACGGTATTCGAATATCACAAGGTGTGTAATCGAGAGCCGCATTTGAATGCGTTCGGCGCGTCGGTCGGCTTCGGCTCGCTCGGTTATTTTACGGACCTGGATGCCGTTACTGGGCGCTTGCCGAGGCTTTTTGTACCGACGACGATCTCGAACCGGCTCTGCGCGCGCGAAAGCGTTCGCGCTCTCGCGCGCTCACCTGTCGGCGACGCTTTTGCTTTGGGTACTTTCAGGACCCGTCTTGAAACACGGACCAAGGAGTCTAACATGTGCGCGAGTCATTGGGTTTTTTTATATTATATTTAAAGCCTAAAGGCGCAATGAAAGTGAAGATACGGCAGGCATTCGTGCCTGAGCCGATCGAGGGAGGATGGCCCGCGTCACGATGCGGGCCCGCACTCCCGGGGCGTCTCGCGCTCATTGCGAGCGGAGGCGCACCCAGAGCGTACACGTTGGGACCCGAAAGATGGTGAACTATGCCTGGTCAGGACGAAGTCAGGGGAAACCCTGATGGAGGTCCGTAGCGATTCTGACGTGCAAATCGATCGTCGGAACTGGGTATAGGGGCGAAAGACTAATCGAACCATCTAG

Sequence found in sample n (length: 308, identities: 300/308 (97.4%), no gap):

CTTGCCGAGGCTTTTTGTACCGACGACGATCTCGAACCGGCTCTGCGCGCGCGAAAGCGTTCGCGCTCTCGCGCGCTCACCTGTCGGCGACGCTTTTGCTTTGGGTACTTTCAGGATCCGTCTTGAAACACGGACCAAGGAGTCTAACATGTGCGCGAGTCATTGGGTTTTTTTATATTATATTTAAAGCCTAAAGGCGCAATGAAAGTGAAGATACGGCAGGCATTCGTGCCTGAGCCGATCGAGGGAGGATGGCCCGCGTCACGATGCGGGCCCGCACTCCCGGGGCGTCTCGCGCTCATTGCGAG

The multiple sequence alignments of the contigs with the reference partial sequence of *I. hookeri* Ixo4 28S ribosomal RNA gene (MH077537.1) are shown in Fig 1. The geolocation of the samples is presented in Fig 2. The figure shows that the sequence of the generated contigs varies from the reference sequence by 8 positions. However, the contigs do not diverge in any position.

**Figure 1.**
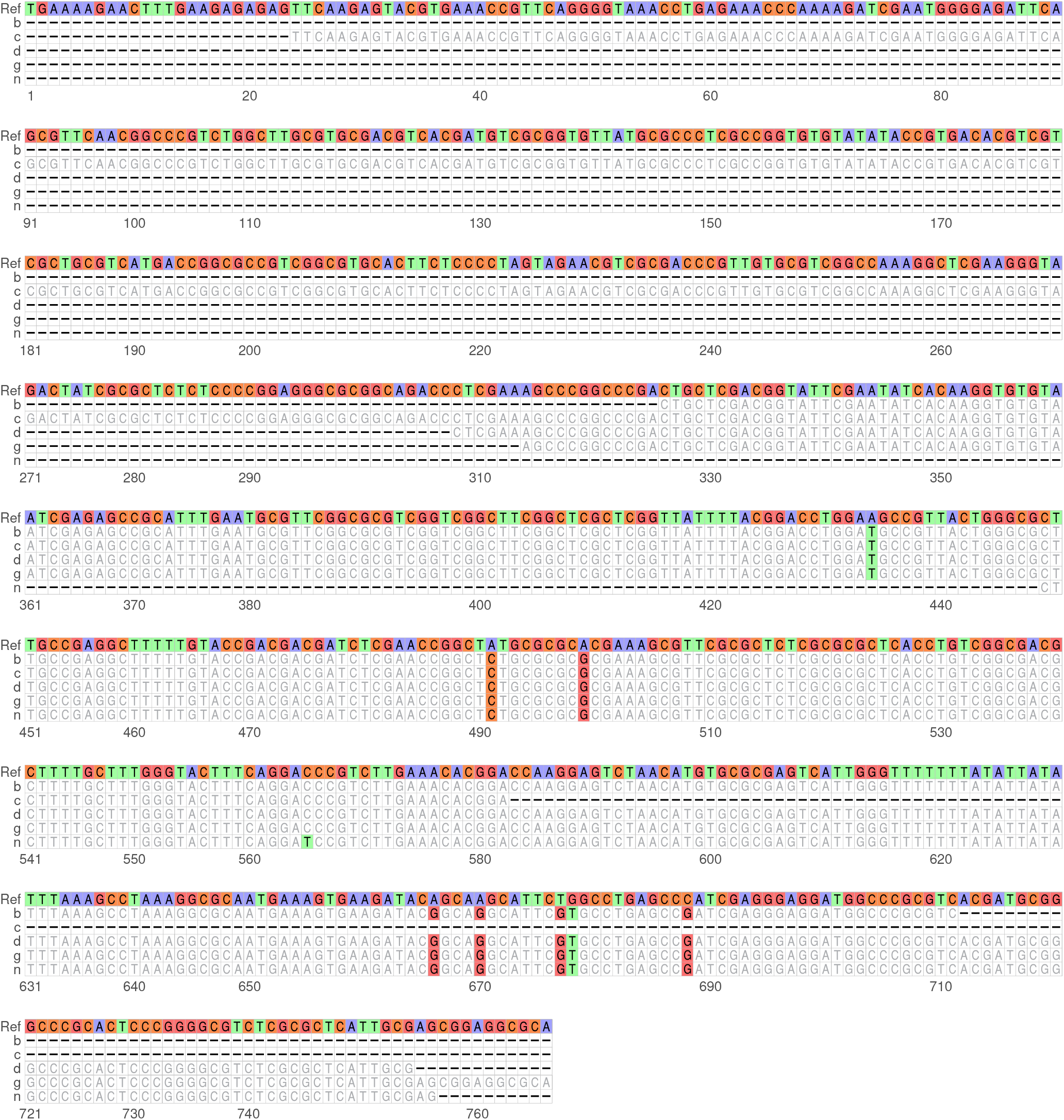
Multiple sequence alignment. The contigs were predicted as origin of *I. hookeri* with the reference partial sequence of *I. hookeri* Ixo4 28S ribosomal RNA gene (MH077537.1). The geolocation of the Hungarian samples are presented in Fig 2.

**Figure 2.**
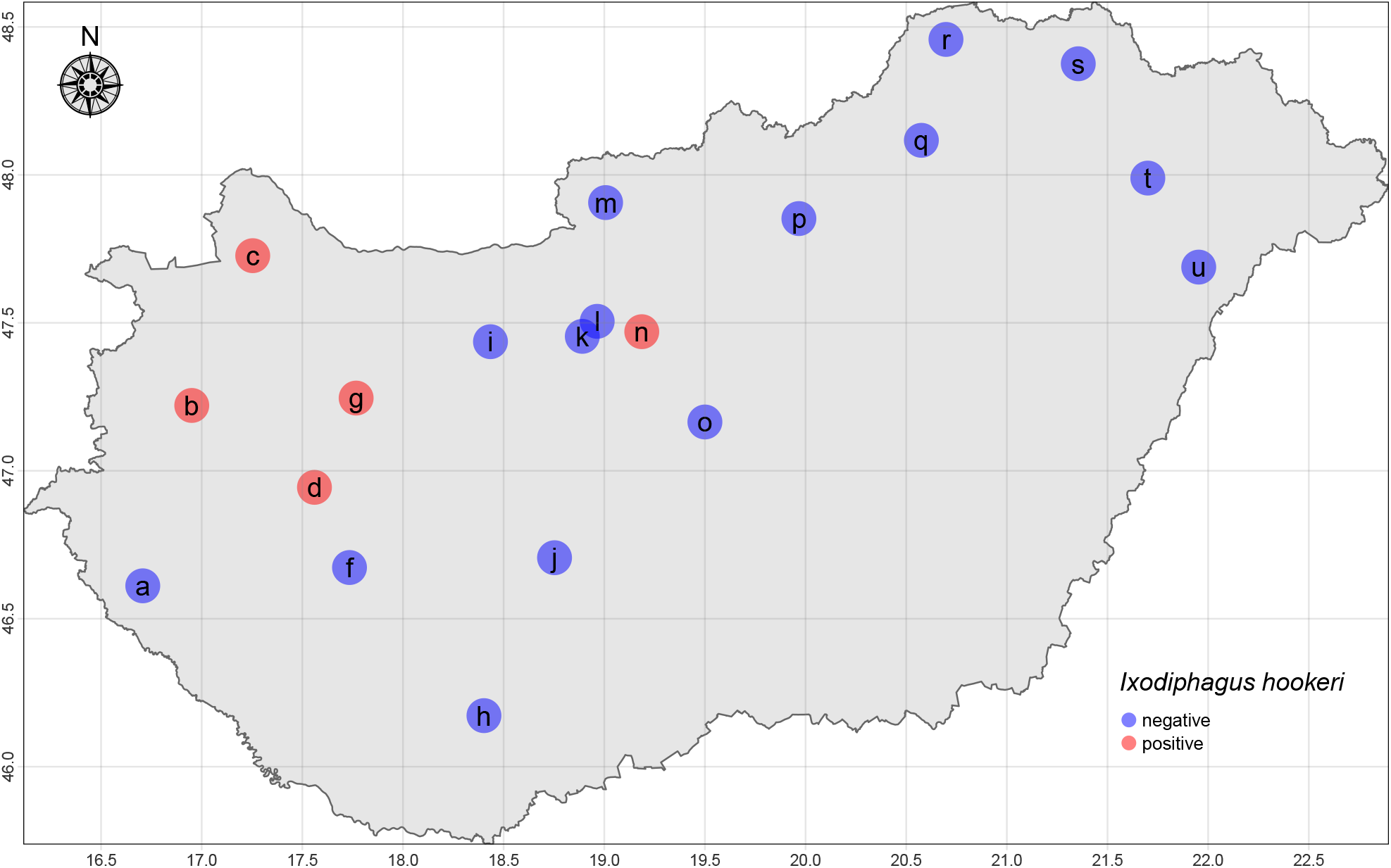
Geopositions of the metagenomicaly analysed samples. The red dots represent the sampling points we have found *I. hookeri* sequences, the blue ones where we have not.

## Discussion

The findings, that no reads deriving from *I. hookeri* were detected in adult *I. ricinus* samples collected between the end of March and the middle of May, while nymphs were associated with this species promotes former theories of the life cycle of the parasitoid wasps.^24,25^ *I. hookeri* eggs may have been laid in the larvae or in nymphs before winter or during spring, as the wasps were formerlry proven to be able to survive harsh winter conditions.^39,40^ Due to the fact that the development of *I. hookeri* can only start after the engorgement of the host and no descriptons of adults living with the parasitoids exsist (thus the survival of infested nymphs is not likely), nymphs collected for our study were either before their first bloodmeal or ingested blood recently.

To our knowledge, this is the first report on evidence of the presence of *Ixodiphagus* wasps, namely *I. hookeri* in Hungary. This finding expands the localities associated with *I. hookeri* within Europe. All except one sampling points that were proven to host *I. hookeri* are located in western Hungary. The cluster of the four western Hungarian sampling points lay close to the borders with Austria and with Slovakia. While the presence of *I. hookeri* has not been published in Austria to our best knowledge, Slovakian reports of the occurrence of the wasps exsist. *I. hookeri* has previously been identified at three locations within Slovakia; near Šoporña, associated to *Haemaphysalis concinna*,^41^ close to the capital of Slovakia, Bratislava, in *I. ricinus*^42,43^ and in the Slovak Karst, isolated from both Ixodes ricinus and *H. concinna*.^40^ According to our hypothesis, the western Hungarian cluster of sampling point b,c,d and g (Sárvár, Mosonmagyaróvár, Sáska, and Pénzesgyőr, respectively) may be associated to the wasp populations described by Bratislava^42,43^ and by Šoporña^41^ in Slovakia. Even though, several geopositions, including sampling point m, p, q, r and s (Nagy-Hideg-Hegy, Kékes, Lillafüred, Aggtelek and Háromhuta respectively) lay relatively close to the Hungarian-Slovakian border, and sampling point r is situated only few kilometers away from the *I. hookeri* associated Slovak Karst,^40^ no tick metagenomes contained read sets aligning to the *I. hookeri* reference 28S ribosomal RNA gene at these sampling points. Shifting slightly to the east, sampling point n (Budapest) represented the closest occurrence of the wasps to Slovakia. Considering the physical proximity among sampling point r and the Slovak Karst where the report of Buczek and colleagues was released, the possibility of receiving false negative results is raised. The basis of receiving false negative results will be described further on.

Other European countries where the presence of *I. hookeri* have been reported include the Czech Republic (detected in the former Czechoslovakia),^44^ Finland,^45^ France,^46,47^ the Georgia,^48^ Germany,^25,27,49^ Italy,^50^ the Netherlands,^28,51^ the United Kingdom,^52^ Russia (detected in the Ussuri forest, in the Asian part of the former Soviet Union)^53^ and Ukraine (detected in the former Soviet Union).^54^ To our knowledge, sampling point d (Sáska) in Hungary represents the southernmost detection point of *I. hookeri* within Central Europe. The detection of *I. hookeri* in Hungary may serve as a novel hint regarding the potential distribution *I. hookeri* at the Balkan Peninsula, where the species appears to be little studied.

As mentioned above, wasp-negative sampling points can actually be wasp-invaded. Even though, next generation sequencing (NGS) based metagenomic approach appears to be just as or even more sensitive as polymerase chain reaction (PCR) based target detection techniques,^29,30^ certain limitations can be addressed. Within the pool of reads deriving from the shotgun sequenced metagenome, that contains genome fragments from every organism present in the sample, lower relative abundance rates of an individual species serve with relatively lower read counts from its genome. In other words, shotgun sequencing preserves the original relative read abundance rates of the various organisms of the samples and may represent less reads of certain species by non-targeted runs.^55,56^ Moreover, the *I. hookeri* reference sequence, that the metagenomic read sets were aligned to only represented a smaller fragment, namely the unique 28S ribosomal RNA gene of the full *I. hookeri* genome. Thus only *I. hookeri* reads deriving from this part of the genome could have been aligned, that further increases the chance of false-negative sampling points for the wasps.

Nevertheless, NGS based approaches have a prospering future within the studies of parasitoids of public health significance, such as *I. hookeri*. According to Collatz and colleagues, large geographical distance and climatic differences (e.g. presence in Africa, Asia, Europe and North America)^25,26,57,58^ may even underlie divergence and distinct taxonomic categorization of *I. hookeri* to different species, subspecies or at least strains.^25^ Concurrently, publications on *I. hookeri* indicate a certain extent of behavioral and feeding variance at different continents.^25,26,59^ In order to assess deeper knowledge of the variances of *hookeri*, or to identify specific traits of subgroups that can be better utilized by the biological control methods, the study of the *I. hookeri* genome, or at least specific genome regions, such as 28S rRNA or 16S rRNA genes may become inevitable, similarly to other weighty insect groups.^51,60,61^

The improvement of our knowledge bank of *I. hookeri* with either traditional or genomic methods could facilitate the assessment of its potential as a means of biological control, while limitations and doubts of the wasps’ biocontrol potential could also be addressed with more expertise. Attempted mass releases of the parasitoid wasps in the U.S and in the former Soviet Union between 1920 and 1940^39,62,63^ were unsuccesful at the noticeable reduction of tick populations. A potential reason for this may be that *I. hookeri* requires high tick host densities and superabundant tick populations to reach its ideal abundance.^59,64^ Inadequate numbers of parasitoids released compared to the geographical areas may have also undermined these trials.^24^ On the other hand, the parasitoid wasps have been transported to the sites of attempted mass releases from great distances, sometimes even from different continents (e.g. from France to the U.S.)^39,62^ without any considerations regarding the host and behavioral preferences of the wasps, that have, since then turned out to be rather specific to their geographic locations of origin.^25,26,59^ Furthermore, we do not know how great the tick populations would be without the endemic *I. hookeri* populations and how much the parasitoid wasps contribute to the maintenance of equilibrium of the communities in which they are included.^65^ Nonetheless, the hypothesis regarding sufficiently high tick host densities and superabundant parasiotid host populations is in line with findings regarding the bioindicative potential of certain insect species, including parasitoid wasps.^66^ If so, this potential may also be worth further observed.

Conclusively, assessment of existing populations and further examinations on entomologic and genomic traits along with ecological roles could help the understanding and exploitation of the *Ixodiphagus* wasps’ potential as a biological tick control method or as a potential bioindicator species.

## Acknowledgements

The project is supported by the European Union’s Horizon 2020 research and innovation program under Grant Agreement No. 874735 (VEO).

## Author contributions statement

NS takes responsibility for the integrity of the data and the accuracy of the data analysis. NS and AGT conceived the concept of the study. EK and MG performed sample collection and procedures. NS participated in the bioinformatic and statistical analysis. AGT and NS participated in the drafting of the manuscript. AGT, NS, and RF carried out the manuscript’s critical revision for important intellectual content. ll authors read and approved the final manuscript.

## Ethics approval and consent to participate

Not applicable.

## Consent for publication

Not applicable.

